# Dynamic Spatial-Temporal Expression Ratio of X Chromosome to Autosomes but Stable Dosage Compensation in Mammals

**DOI:** 10.1101/2021.08.11.455930

**Authors:** Sheng Hu Qian, Yu-Li Xiong, Lu Chen, Ying-Jie Geng, Xiao-Man Tang, Zhen-Xia Chen

## Abstract

In the evolutionary model of dosage compensation, per-allele expression level of the X chromosome was proposed to have two-fold upregulation, compensating for its dose reduction in males (XY) compared to females (XX). However, the upregulation of X chromosome is still in dispute, and comprehensive evaluations are still lacking. By integrating multi-omics datasets in mammals, we investigated the expression ratios and underlying pattern of X to autosomes (X:AA ratio) and X to orthologs (X:<u>XX</u> ratio) at the transcriptome, translatome, and proteome layers. The results indicated a dynamic spatial-temporal X:AA ratio during development in human and mouse. Meanwhile, by tracing the evolution of orthologous gene expressions in chicken, platypus, and opossum, we found a constant expression ratio between X-linked genes in human and their autosomal orthologs in other species (X:<u>XX</u> ~1) across tissues and developmental stages, demonstrating stable dosage compensation in mammals. We also revealed that different epigenetic regulations could shape the higher tissue- and stage-specificity of X-linked gene expression, and affect X:AA ratios. We conclude that the dynamics of X:AA ratios are attributed to the different gene contents and expression preferences of the X chromosome, instead of the stable dosage compensation.

## Introduction

The therian X and Y chromosomes originated from a pair of ancestral autosomes about 190–166 million years ago (MYA) [1]. Evolutionary degeneration of the Y chromosome caused dose reduction of X-linked genes [2, 3]. The dosage compensation of sex chromosome and its underlying mechanisms attracted long-term attention in molecular evolution [4]. Susumo Ohno (1967) proposed that the expression of X-linked gene should be doubled to compensate for the dose reduction of X chromosome in males (XY), followed by the inactivation of one X chromosome in females (XX) to avoid X-tetrasomy formation in female cells and to equalize gene expression level between sexes [5]. Ohno’s hypothesis lays the theoretical foundation for the evolution of sex chromosome dosage compensation [1, 6].

Several previous studies have reported the X upregulation in some certain mammalian tissues (X:AA ratio ~1) by microarray [7–9], RNA sequencing [10–13], and recently used ribosome sequencing [14], but Ohno’s hypothesis has also been challenged [15, 16]. Due to the unavailability of the ancestral proto-X (<u>X</u>), the above-mentioned tests of Ohno’s hypothesis depended on indirect calculation of the expression ratio of present-day X to autosomes (X:AA). An innovative study directly compared the human X-linked genes with their orthologs in chicken [17], an outgroup species diverged from therians (~310 MYA) prior to the origin of X chromosome, but its inclusion of unexpressed genes in the evaluation of dosage compensation is disputable [10]. In addition, in the existing studies, only limited tissues and certain developmental stages have been investigated, which might be nonrepresentative [18, 19]. Fortunately, the emergence of developmental transcriptome data covering multiple tissues, developmental stages and species enables us to perform a comprehensive analysis of the regulation and evolution of dosage compensation [20, 21]. We thus took advantage of the multi-omics datasets, and performed a comprehensive analysis on the X:AA ratios and X:<u>XX</u> ratios to test Ohno’s hypothesis. Our analysis reveals per-allele upregulation of X chromosome at the transcriptome, translatome and proteome layers across tissues and developmental stages in mammalian species, systematically validating Ohno’s hypothesis of dosage compensation in mammals.

## Results

### Tissue-dependent X:AA expression ratios in mammals

The recent release of RNA-seq data across tissues of multiple mammalian species makes possible a comprehensive examination of Ohno’s hypothesis with X:AA ratio. We used public RNA-seq data of 32 main healthy adult human tissues [22], and found tissue-dependent X:AA ratios, ranging from 0.12 (pancreas) to 1.4 (cerebral cortex) (**Figure 1A**). The majority of X:AA ratios of our investigated tissues fitted Ohno’s hypothesis with X:AA ~1, except for pancreas (0.12), saliva-secreting gland (0.45), skeletal muscle tissue (0.51), and liver (0.51). We also applied Genotype Tissue Expression (GTEx) to validate the results [23]. The lowest X:AA ratio (0.24) and the highest ratio (1.27) were still observed in human pancreas and neural tissues, respectively (Figure S1), indicating great consistency between different datasets. We further estimated tissue-wide X:AA ratios in other mammals, including rhesus (*Macaca mulatta*), mouse (*Mus musculus*), rat (*Rattus norvegicus*), and rabbit (*Oryctolagus cuniculus*) (Figure 1B, Figure S2, methods) [20, 21, 23, 24]. Among the shared investigated tissues across these five species, we observed the consistent pattern that the highest ratio existed in brain and the lowest ratio existed in liver, demonstrating tissue-dependent X:AA ratios at the transcriptome layer.

**Figure 1.**
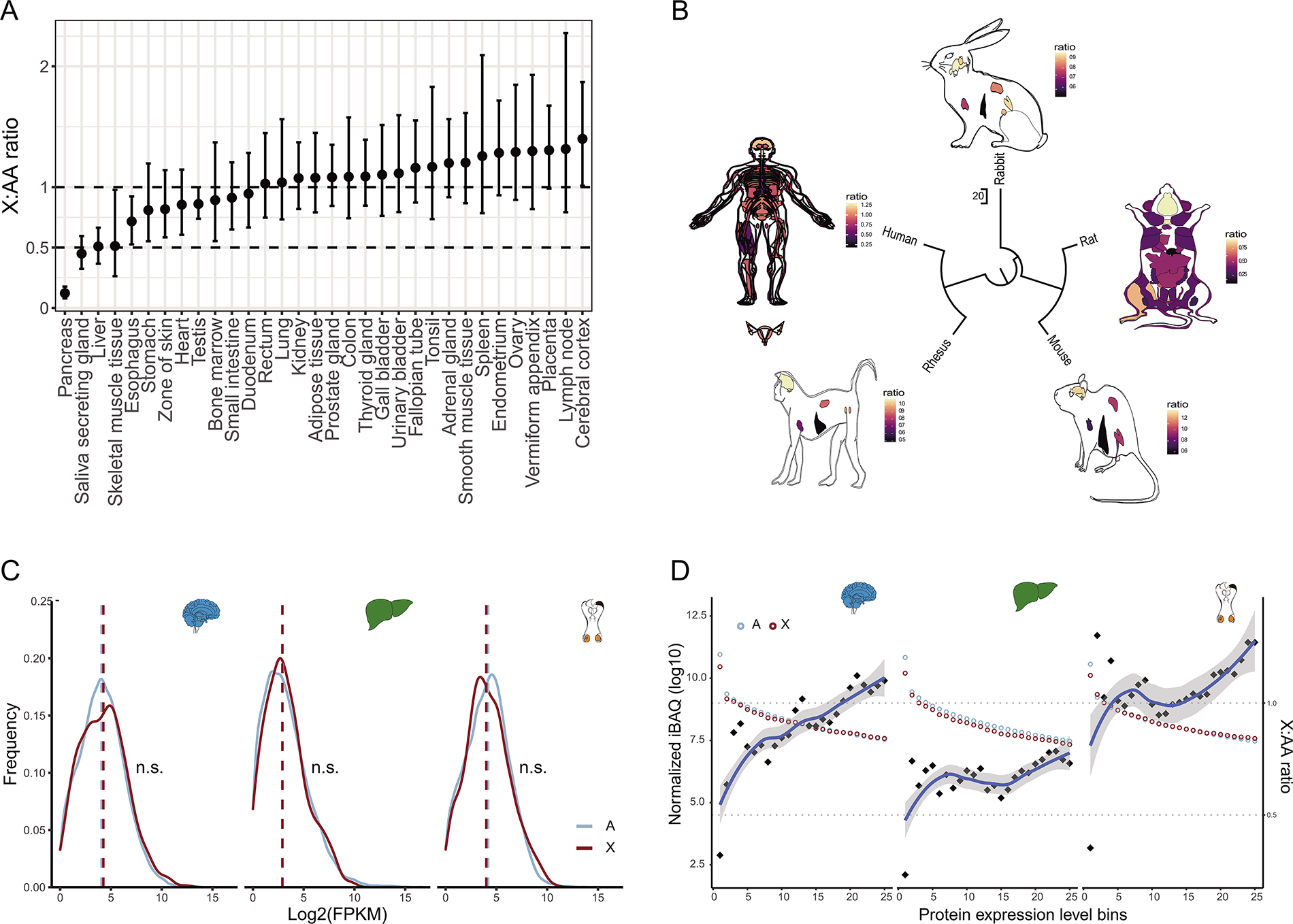
Expression ratio of X:AA across mammalian anatomies. **A**) X:AA ratio across 32 adult human tissues. Black points indicate mean value, and the error bars represent 90% confidence interval. **B**) X:AA ratio in human (50 tissues), rhesus (*Macaca mulatta*, 7 tissues), mouse (*Mus musculus*, 22 tissues), rat (*Rattus norvegicus*, 7 tissues), and rabbit (*Oryctolagus cuniculus*,7 tissues). Human female reproductive tissues, including ovary, endometrium, vagina, fallopian tube, uterus, uterine cervix, amd ectocervix, were attached. **C**) Expression distributions of X-linked and autosomal genes in human brain, liver, and testis at the translatome layer. **D**) Comparison of protein abundance between X-linked genes and autosomal genes in human brain, liver, and testis. The protein abundances of X-linked and autosomal genes are separately divided into 100 expression bins, and top 25 bins are used for analysis.

Given that mRNAs are functionally and phenotypically relevant only after they are translated into protein and the expression of protein-coding genes may frequently be regulated at multiple layers after transcription [25–27], so transcriptome might offer an incomplete picture of gene activity. Therefore, the protein abundance would be better to represent the dosage compensation, and the X:AA might be closer to the actual ratio at the translatome layer than at the transcriptome layer. To achieve a more precise evaluation of X:AA ratio, we took advantage of ribo-seq data [14], which allowed the direct measurement of translation [28]. We found the distribution of X-linked genes at the translatome layer greatly resembled that of autosomal ones in brain (X:AA = 1.13), testis (X:AA = 0.88), and even in liver (X:AA = 0.97) (Figure 1C, Figure S3) (P > 0.05, wilcoxon test). Notably, in liver, no upregulation of X chromosome has been detected at the transcriptome layer using multiple RNA-seq datasets in human and mouse (Figure 1A). We speculated that X:AA ratio in a certain tissue deviated from 1 at the transcriptome layer could restore to 1 at other gene regulatory layers due to extensive buffering between different expression layers [14, 29, 30], and vice versa.

We further incorporated mass spectrometry proteome data across various tissues, which would allow a more direct measurement of protein abundance, to verify the results from ribo-seq data [22]. Considering the current proteomic coverage and resolution, X-linked genes and autosomal genes were separately divided into 100 bins in terms of normalized intensity-based absolute quantification (iBAQ), and only the first 25 X and autosomal bins were investigated. The X:AA ratio of protein concentrations between matched bins were approximately 0.75 across all tissues and reached 1 in brain, endometrium, placenta, and testis (Figure 1D, Figure S4). Interestingly, we also noticed that in pancreas, the X:AA ratio was about 1 at the proteome layer, while the lowest X:AA ratio was observed at the transcriptome layer from different datasets, and such inconsistency might be attributed to poor correlation between transcriptome and proteome data [22]. Most of the tissues (55%) concordantly showed X:AA≈ 1 at the proteome layer (including adrenal gland, appendix, brain, colon, endometrium, esophagus, fallopian tube, heart, lung, pancreas, placenta, prostate, smooth muscle, spleen, testis, and thyroid), despite incomplete upregulation in some tissues, which reconfirmed the upregulated expression of X chromosome.

### Dynamic developmental X:AA ratios in mammals

To determine whether X:AA was dependent on development stages, we measured X:AA ratio in different development stages from early organogenesis to adulthood across seven major tissues (brain, cerebellum, heart, kidney, liver, ovary, and testis) in human and mouse with the recently released RNA-seq dataset [20]. To avoid possible biased evaluation of the X:AA ratio since “filter-by-expression” strategy was only limited to complete dosage compensation model [31], we selected a series of expression cutoffs (FPKM from 0-1) with an additional group retaining all genes as the control. Interestingly, the X:AA ratio showed developmental dynamics in a tissue-specific manner in both human and mouse. In neural tissues (brain and cerebellum), we found a noticeable positive correlation between the X:AA ratio and development stage (rho = 0.86, P = 2.49×10^−7^) (**Figure 2A**, Figure S5-6).This correlation was insensitive to different expression thresholds. Conversely, in the remaining tissues, X:AA ratio was negatively correlated with development stage (Figure 2B, C). In addition, in liver, X:AA ratio was close to 1 at early development stage and decreased to 0.5 with development in human (Spearman correlation rho = −0.51, p = 0.012) and mouse (rho = −0.96, p = 0) (Figure 2B, Figure S6E). As for reproductive tissues, the human ovary samples only covered the developmental stages from 4 weeks post-conception (WPC) to 18 WPC. Among these stages, the X:AA ratio in testis and ovary was similar, ranging from 1 to 1.5 (Figure 2C-D, Figure S6F). While X:AA ratio dropped to 0.5 after reproductive maturity, presumably due to the X-hypoexpression of numerous germ cells after meiotic sex chromosome inactivation. Consistently, one previous study has reported that low X:AA ratio might be essential for male sperm cell development [32]. In mouse, the X:AA ratio in testis was comparable to that of in ovary (about 1) before 2 weeks post-born (WPB) and dropped to 0.5 at 4 WPB and 9 WPB, while in ovary at 4 WPB and 9 WPB, the X:AA ratio was still near 1. One past work observed upregulation of X-linked gene in specific cell type, mouse oocyte, at immature stage instead of mature stage [13], which showed a developmental stage-dependent X:AA ratio. The ovary tissue contained many cell types, including germ cells, immune-related cells, and other somatic cells [33]. Although the oocyte harbored two active X chromosomes and exhibited a low X:AA ratio (<1) at mature stage [13], the X:AA ratio in mature ovary was near 1 in our data, which might be that the X:AA ratio in ovary was mixed and reflected the average level of all cell types. Since the testis was the most distinct tissue than other tissues including ovary, such as cell composition and chromatin state, the developmental pattern of X:AA ratio in testis might not be identical to any other tissues.

**Figure 2.**
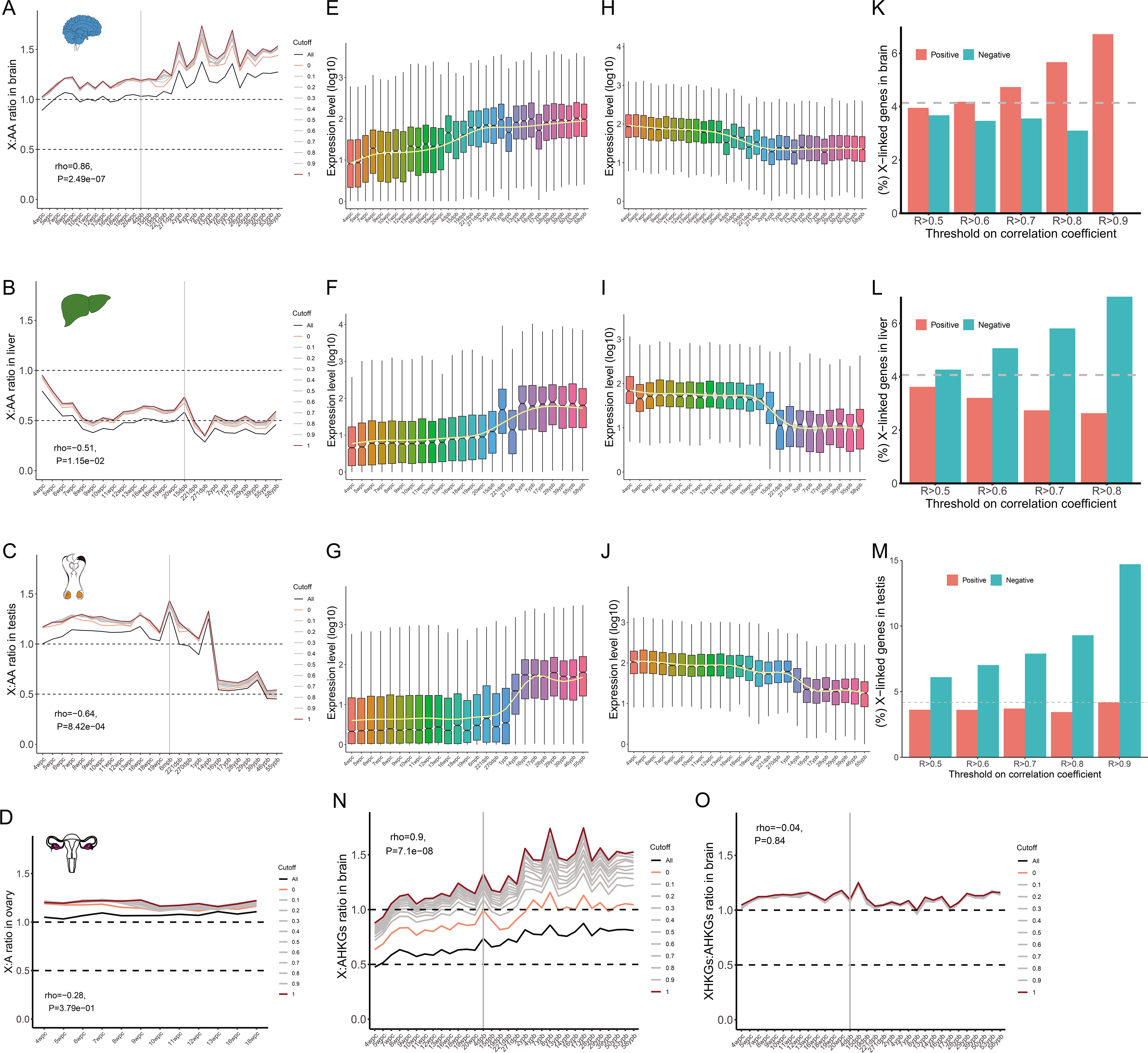
Dynamics of X:AA ratio during development. **A**) X:AA ratio in brain, **B**) in liver, **C**) in testis, and **D**) in ovary. **E**) Expression profile of stage-positively-correlated genes in brain (rho > 0.8), **F**) in liver, and **G**) in testis. **H**) Expression profile of stage-negatively-correlated genes in brain (rho < −0.8), **I**) in liver, and **J**) in testis. **K**) Percentage of X-linked stage-correlated (positive and negative) genes under different correlation thresholds in brain, **L**) in liver, and **M**) in testis. R > 0.5 indicates that the absolute spearman correlation coefficients (rho) are higher than 0.5. Red color represents stage-positively-correlated genes, while blue color denotes stage-negatively-correlated genes. **N**) Expression ratio of X-linked genes to autosomal housekeeping genes, AHKGs. **O**) Expression ratio of X-linked housekeeping genes, XHKGs, to AHKGs

In this study, we also found that the X chromosome was doubly upregulated (X:AA ratio = 1) at early stage across all tissues, suggesting strong similarity in development programs at the earliest stage (4 weeks post-conception) among different tissues and increasing molecular and morphological difference of tissues during development [20, 34]. Therefore, development stage should be taken into account when we examined the expression of X-linked genes, since opposite conclusions that dosage compensation was accepted or rejected could be reached, even if the same tissue but in different development stages was investigated.

To reveal the reason for the dynamics of X:AA ratio, we identified the genes whose expressions were correlated with development stage (Spearman correlation, rho > 0 for positive correlation, rho< 0 for negative correlation) using a previously described method [21]. The expression levels of stage-positively-correlated genes (rho > 0.8, P < 0.05) and stage-negatively-correlated genes (rho < −0.8, P < 0.05) in brain, liver, and testis were presented in Figure 2D-I. GO analysis showed that these stage-correlated genes were involved in biological processes of corresponding tissues. In brain, the stage-positively-correlated genes were enriched in vesicle-mediated neurotransmitter transport and synapse signal release pathways; in liver, they were enriched in fatty acid and organic acid metabolic pathways; and in testis, they were enriched in spermatid development, differentiation, and fertilization pathways (see Supplemental Table S1), implying the function demand and expression timing during development led to the dynamic X:AA ratio. We also applied other correlation thresholds (from 0.5 to 0.9) to screen stage-correlated genes. In brain, as the thresholds became more stringent, more stage-positively-correlated genes were located on the X chromosome than on autosomes, but stage-negatively-correlated genes exhibited an opposite pattern, (Figure 2J), suggesting that more X-linked stage-positively-correlated genes and less stage-negatively-correlated genes might result in higher X:AA ratio during development. The above speculation was confirmed by the evidence that the less stage-positively-correlated genes and more stage-negatively-correlated genes were enriched on X chromosome in liver and testis (Figure 2K, L). Our results revealed that development stage-correlated genes contributed to the dynamics of X:AA ratio.

To further analyze the expression dynamic of X-linked genes, we calculated the expression ratio between X-linked genes and autosomal housekeeping genes (AHKG). Since the housekeeping genes are constitutively and steadily expressed, the ratio of X-linked genes to AHKGs would follow the expression pattern of X-linked genes. Consistent with our expectations, the ratio of X-linked genes: AHKGs in brain was still positively correlated with development stages (Figure 2N), suggesting the increased expression of X-linked genes during development. When we compared the X-linked housekeeping genes (XHKGs) with AHKGs, the XHKGs:AHKGs ratio changed to be constant, instead of dynamic (Figure 2O), confirming again that the above pattern was attributed to stage-correlated genes.

### Expression maintenance of X-linked genes in mammalian evolution

As Ohno has stressed, the expression of the present-day X chromosome should be doubled (X:<u>XX</u> ~1) to maintain the same expression level with ancestral autosomes that have evolved into sex chromosomes (proto-X/Y, proto-X hereafter marked as <u>XX</u>). Ohno’s hypothesis of X:<u>XX</u> ~1 would be equivalent to X:AA ~1 under two assumptions: 1) gene expression on the present autosomes (AA) is comparable to that on the proto-autosomes (<u>AA</u>) (AA:<u>AA</u> ~1); 2) gene expression on the proto-X is comparable to that on the proto-autosomes (<u>XX:AA</u> ~1) [17]. To directly test dosage compensation and determine whether the dynamic X:AA ratio was responsible for the dynamic dosage compensation, we investigated the expression ratio of X-linked genes in human (X) to the autosomal orthologous genes (<u>XX</u>, 1:1 orthologs) in opossum, platypus, and chicken (X:<u>XX</u>) (**Figure 3A**, see methods) [17]. The phylogenetic distance of the three species from human was as follows: opossum < platypus < chicken. The expression level of opossum <u>XX</u> (R = 0.5-0.67 in transcription, and 0.3-0.67 in translation) was more similar to that of human X than platypus (R = 0.45-0.53 in transcription, and 0.3-0.71 in translation) and chicken (R = 0.11-0.46 in transcription, and 0.17-0.46 in translation) (Figure S7-8), suggesting the expression of autosomal orthologs on the closer relatives could better represent that of human X-linked genes. At the transcriptome layer, the X:<u>XX</u> ratio was 0.57 for human:chicken on average (0.66 in brain, 0.57 in liver, and 0.49 in testis), and 0.62 for human:platypus (0.61 in brain, 0.74 in liver, 0.5 in testis), and 0.78 for human:opossum (0.82 in brain, 0.77 in liver, 0.75 in testis) (Figure 3B-D). The comparison of human with opossum, the closest relative of human among the three species, resulted in the X:<u>XX</u> ratio closest to 1. For all the three species, X:<u>XX</u> ratio was closer to 1 at the translatome layer than at the transcriptome layer across all the three tissues (Chicken, 0.83, 0.78, and 0.85; Platypus, 0.82, 0.77, and 0.90; Opossum, 0.92, 0.86, and 0.92, in brain, liver, and testis, respectively) (Figure 3E-G). These observations indicated that the current X chromosome was upregulated about two folds to compensate for the decay of Y chromosome in a tissue-independent manner, and that the dynamic X:AA ratios across tissues was not due to dosage compensation.

**Figure 3.**
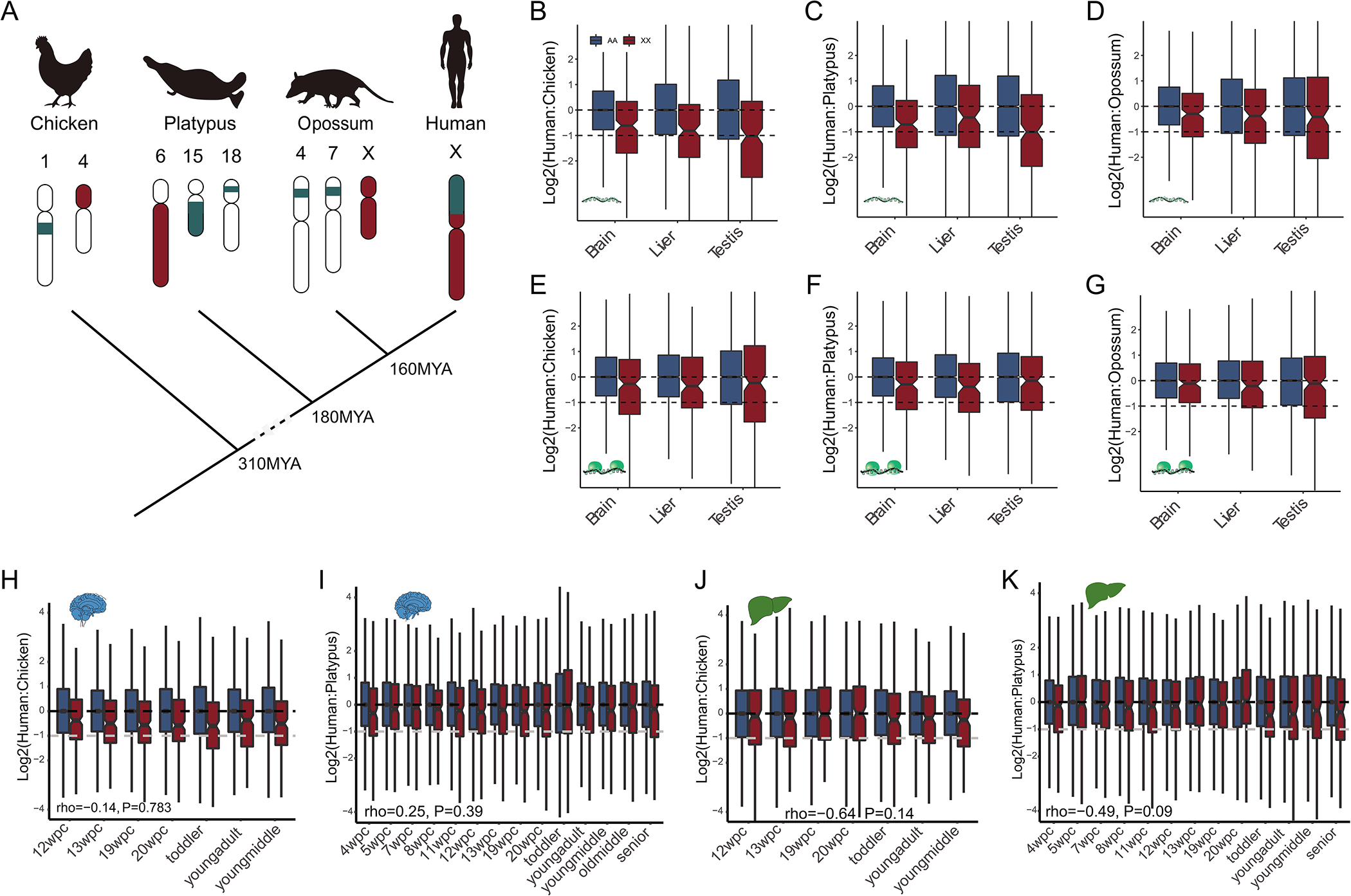
Comparison of expression levels of orthologous genes between human and outgroup species (chicken, platypus, and opossum). **A**) Origin and evolution of mammalian X chromosome. The genes on the human X chromosome are located on autosomes in chicken (birds, on chromosome 1 and 4) and platypus (monotremes, on chromosome 6, 15, and 18). The X chromosome of opossum (marsupials) stands for the most ancient segment of the mammalian X chromosome (shown in red). **B**) Human X:chicken <u>XX</u> ratios in brain, liver, and testis after the median of human AA:chicken <u>AA</u> ratios is normalized into 1 at the transcriptome layer, **C**) platypus as outgroup species, **D**) opossum as outgroup species. **E**) X:<u>XX</u> ratio at the translatome layer with chicken as outgroup species, **F**) platypus as outgroup species, **G**) opossum as outgroup species. **H**) Human X:chicken <u>XX</u> ratios at Carnegie matched stages at the transcriptome layer in brain. **I**) Human X:platypus <u>XX</u> expression ratios at Carnegie matched stages at the transcriptome layer in brain. **J**) Human X:chicken <u>XX</u> expression ratios at Carnegie matched stages at the transcriptome layer in liver. **K**) Human X:platypus <u>XX</u> expression ratios at Carnegie matched stages at the transcriptome layer in liver.

Using chicken and opossum <u>XX</u> as control, we further addressed whether the developmental dynamics of current X chromosome was conferred by dosage compensation based on X:<u>XX</u> ratio at matched Carnegie stages [20]. Unlike the X:AA ratio, the X:<u>XX</u> ratio was not stage-correlated (Figure 3H-K, Figure S9), suggesting again that the dynamic X:AA ratios across developmental stages were not resulted from dosage compensation.

### Higher tissue- and stage-specificity of gene expression on the X chromosome may be responsible for the dynamic X:AA ratios

The inconsistency in dynamics between X:<u>XX</u> ratio and X:AA ratio could have resulted from the different gene contents and expression preferences of the X chromosome due to its monosomy [35–37]. We compared the expression patterns of X-linked and autosomal genes in mammals using RNA-seq data across seven tissues (brain, cerebellum, heart, kidney, liver, ovary, and testis) and two dozen developmental stages [20]. We found that about 34% of all X-linked genes showed higher expression level in testis than in other tissues in human and mouse, followed by brain and cerebellum (**Figure 4A**), which was consistent with previous findings [38, 39]. Such tendency was robust when the extended 32 tissues were investigated, and the X chromosome favored testis with 26.4% genes significantly enriched (fisher exact test P = 6×10^−7^) (Figure 4B). These RNA sequencing results were in accordance with those from mass spectrometry (Figure S10A). The expression of X-linked genes exhibited higher tissue- and developmental stage-specificity than that of autosomal genes across all tissues in human and mouse (Figure 4C,D, Figure S10B,C). Similar phenomenon was also observed in rhesus, rabbit, rat, and opossum (Figure S11-S12), demonstrating the preference of dynamic expression of X chromosome in mammals.

**Figure 4.**
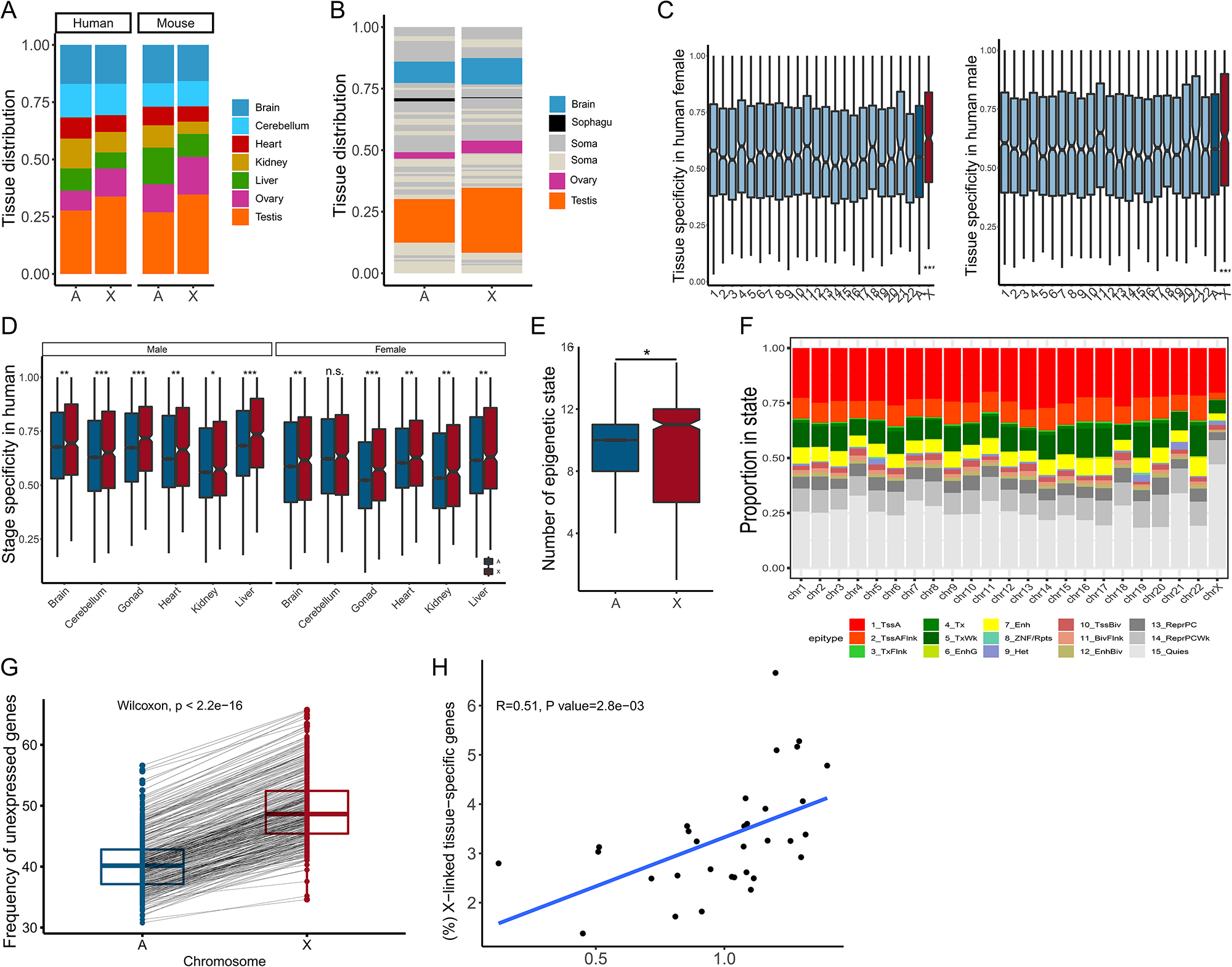
Expression pattern of the X chromosome and autosomes. **A**) Tissue distribution in which genes showed maximum expression at transcriptome layer. **B**) Distribution of the extended 32 tissues in which genes showed maximum expression at transcriptome layer. **C**) Tissue specificity of genes across all chromosomes. A theoretical box line (blue) generated by averaging tau values of autosomal genes. **D**) Developmental stage-specificity of genes expression. **E**) Numbers of epigenetic states within each gene. **F**) Percentages of bases within genes on each chromosome annotated with each epigenetic state, sum of all epigenomes. **G**) Percentages of unexpressed genes (FPKM < 1). Each line indicates the two linked points in the same tissue. **H**) Correlation between the X:AA ratio and percentage of X-linked tissue-specific genes, and a point indicates a tissue.

To explain for the higher tissue- and developmental stage-specificity of X chromosome expression, we quantitatively investigated the epigenetic modifications contributing to selective expression of genes across human tissues using Roadmap Epigenomics Project data [40, 41]. We analyzed the dynamics of each promoter to obtain their epigenetic profile. The results indicated the median of epigenetic states displayed by X-linked genes was 11 across all epigenomes, which was higher than that by autosomal ones (with median of 10) (P < 0.05, Wilcoxon test), supporting that X chromosome acquired variable gene regulation (Figure 4E). We further calculated the total proportion of promoters annotated with each epigenetic state across all Roadmap epigenomics. Half of X-linked gene promoters were in the quiescent state (15_Quies) (Figure 4F, Figure S13) which represented the lack of five constituent histone modifications (H3K4me3, H3K4me1, H3K36me3, H3K9me3, and H3K27me3), followed by active TSS state and regulatory chromHMM state (1_TssA, 2_TssAFlnk, 3_TxFlnk, 6_EnhG, 7_Enh). More X-linked genes in quiescent state than autosomal ones suggested an enrichment of more unexpressed genes. As expected, X chromosome had significantly higher percentage of unexpressed genes (FPKM < 1) than autosomes in human (~1.22 times, P < 3.19×10^−3^) and mouse (~1.26 times, P < 3.15×10^−3^) (Figure 4G, Figure S14A). The higher percentage of unexpressed genes was enriched on X chromosome than on autosomes when a less strict cutoff (FPKM = 0) was used in human (~1.7 times) and mouse (~1.58 times) (Figure S14B, C). If we examined further this percentage by tissue type, we still observed the higher percentage of unexpressed X-linked genes than autosomal genes (Figure S15-18). Specifically, X chromosome exhibited highest percentage of unexpressed genes in liver (58% in human, 64% in mouse). The lowest percentage of unexpressed genes was found in gonad and brain, which was in line with the previous reports on brain and testis preference of X chromosome [36, 38]. We also noted that the percentage of unexpressed genes in ovary was as low as in testis, which agreed with previous results that the genes related to spermatogenesis and oogenesis were enriched on the X chromosome [32, 42]. Since the expressed genes should be compensated, these genes with significant expression difference in tissue- and stage-specificity beween X chromosome and autosome should be stressed when we tested Ohno’s hypothesis.

We further checked whether the dynamic X:AA ratio resulted from the different tissue-/stage-specificity of gene expression between X chromosome and autosomes. As mentioned above, 32 human tissues showed the varied X:AA ratios. We identified approximately 290 (in pancreas) to 4180 (in testis) tissues-specific genes from 32 human tissues (see Methods Section). The X:AA ratio in a certain tissue was positively correlated with the percentage of X-linked tissue-specific genes in the corresponding tissue (Figure 4H). After further dividing the tissues into X:AA~0.5 group (pancreas, saliva secreting gland, liver, and skeletal muscle tissue) and X:AA~1 group (the rest 28 tissues from 32 tissues), we found that tissues in X:AA~1 group had more tissue-specific genes (Figure S19), validating that X-linked tissue-specific genes contributed to the X:AA ratio in corresponding tissue.

To illustrate the spatiotemporal expression specificity of X-linked genes in detail and raise more biological evidence, two X-linked genes *F9* and *SPANXD* were taken as examples. *F9* encodes vitamin K-dependent coagulation factor IX in liver, plays role in the blood coagulation cascade, and mutation in *F9* would lead to hemophilia B or Christmas disease. Cross-species developmental transcriptome data showed that *F9* was exclusively expressed in liver and exhibited increased expression level during development, which was conserved in tetrapods (**Figure 5A**, Figure S20). To validate expression specificity of *F9*, we searched this gene in NCBI and ARCHS4 which collected more than 100,000 human RNA-seq samples [43]. Among the 72 tissues across 9 systems of the human body, *F9* was biasedly expressed in the liver-associated cells, including liver, hepatocyte, and kupffer cell (Figure 5B, Figure S21). Additionally, *SPANXD* encodes testis-specific proteins, which regulate expression of some testis-specific genes necessary for the morphological changes of male germ cell and required for the formation of mature spermatozoa. We found that *SPANXD* was solely expressed in testis in multiple large-scale RNA-seq datasets (Figure 5B, Figure S21). Moreover, *SPANXD* was silenced at embryo and young age, and started to be expressed at adolescent age (13-14y) (Figure 5C), confirming the consistency between its function and expression timing.

**Figure 5.**
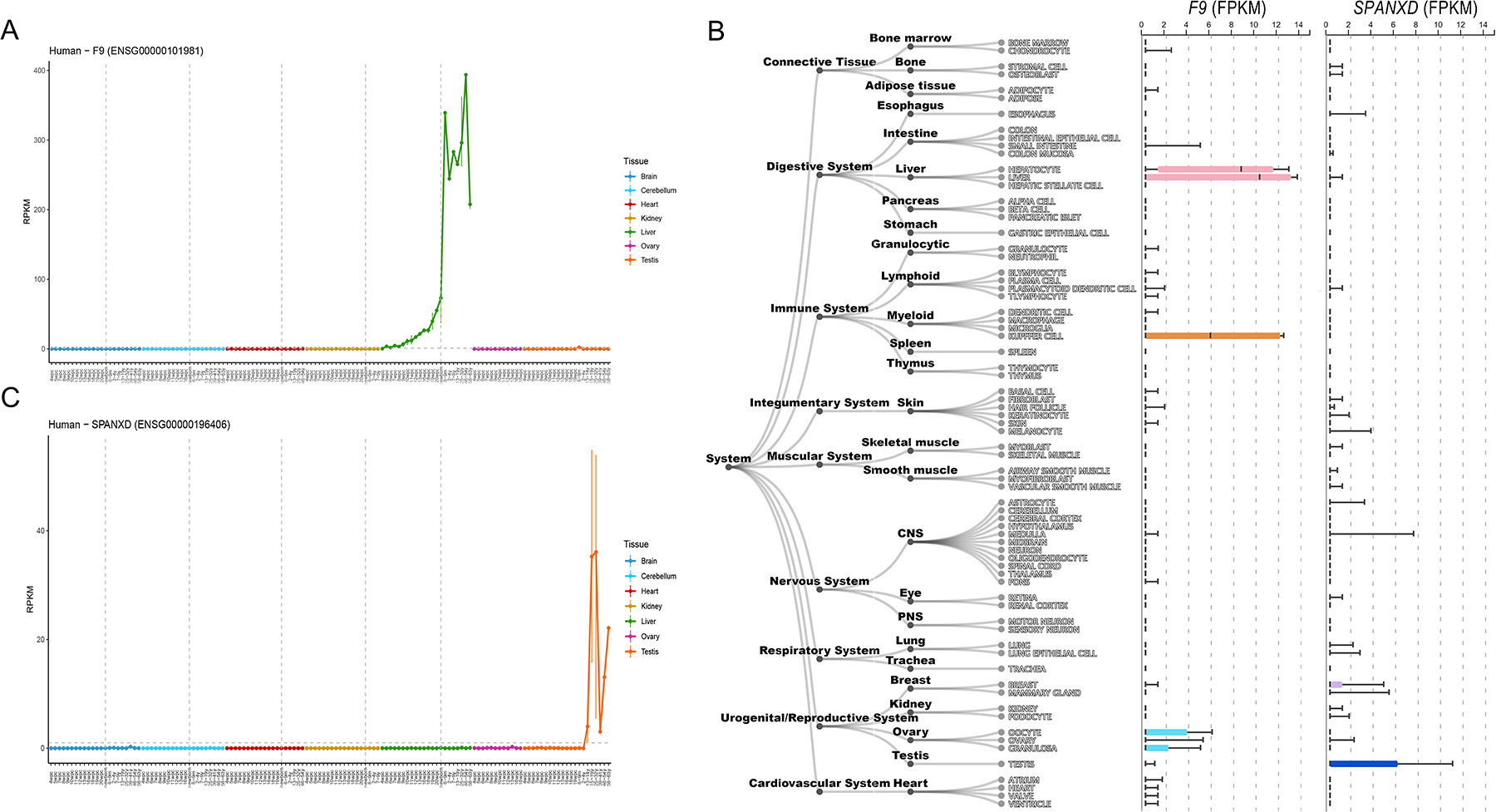
Expression level of *F9* and *SPANXD*. **A**) Expression level of *F9* during development in human across seven tissues. **B**) Expression level of *F9* and *SPANXD* across nine human systems from ARCHS4. **C**) Expression level of *SPANXD* during development in human across seven tissues.

## Discussion

As Ohno hypothesized, therian X-linked genes should be upregulated two folds to counteract the degeneration of Y homologs and to reach the same expression levels of ancestral X-linked genes [6]. Because of the unavailability of ancestral X chromosome, Ohno’s hypothesis is often indirectly tested by comparing the gene expression level between X and AA under the assumption that the gene expression levels are comparable among <u>XX</u>, <u>AA</u>, and AA. Since the first empirical test was conducted, however, there has been an ongoing debate on the validity of Ohno’s hypothesis for fifteen years [7, 9–12, 14–17, 44–46]. These previous studies tended to be limited to RNA sequencing or only focus on a subset of tissue expression data. In this study, in combination with GTEx and ENCODE project data [23, 24], we dramatically expanded the sequencing datasets [22] and provided a comprehensive profile of X:AA ratio across tissues, developmental stages, and species. Moreover, despite the random errors of RNA-seq and the poor correlation between mRNA concentration and protein abundance [22], we still found the comparable gene expression levels between X chromosome and autosomes at translatome and proteome layers, suggesting the upregulation of current X chromosome at three expression layers. It should be noted that previous studies did not detect dosage compensation at the proteome layer [17, 46], which might be due to the limited resolution of mass spectrometry [14] and the difference in cell biology between cell line and tissue.

Another limitation of previous works lies in that merely high-throughput data of adult tissues have been investigated, while the data of adult even aged tissues alone could not provide a full picture of X:AA ratio across the entire life span. Additionally, since all tissues are developed from a single zygote, but it is hard to determine when and how different X:AA ratios are produced in these adult tissues. To fill this gap, we applied the published developmental transcriptome data ranging from early organogenesis to adulthood, and focused on seven tissues representing three layers: liver (endoderm), kidney, heart, ovary, and testis (mesoderm), brain and cerebellum (ectoderm) in human and mouse. Our results revealed a dynamic X: AA ratio during development, and the difference in chromosomal distribution of stage-correlated genes contributed to the diverse dynamics of X:AA ratio among three germ layers. We also noted that the X chromosome was fully upregulated at the earliest development stage across seven tissues, but the X:AA ratio gradually differed during development, demonstrating the high similarity of expression programs at early stage and increasing discrepancy during development [20]. One limitation of our study is that only bulk RNA-seq data we used to examine Ohno’s hypothesis, although multiple tissues and development stages covered in bulk RNA-seq enable us comprehensively profile dynamic pattern of X:AA ratio, recently emerging single-cell technology provides new insight into the transcriptional status at single-cell resolution [47]. In the future, the upregulation of X chromosome and the heterogeneity degree of X:AA ratio could be investigated at single-cell level.

The above-mentioned tests of Ohno’s hypothesis based on X:AA ratio are indirect. To directly examine dosage compensation, we calculated X:<u>XX</u> ratio at the transcriptome and translatome layers. Due to the extensive buffering between the expression levels, the expression difference between species is smaller at the translatome layer than at the transcriptome layer [14], thus resulting in a more robust evaluation of dosage compensation. Furthermore, the Carnegie matched stages comparison of human and chicken or opossum showed a constant X:<u>XX</u> ratio value close to 1, in contrast with the dynamic pattern of X:AA ratio. One previous study claimed that gene expression of X chromosome was halved, thereby refuting Ohno’s hypothesis [17]. Repeating their analysis using the same data, we found that the human X chromosome harbored more unexpressed genes than human autosomes, chicken <u>XX</u>, and chicken <u>AA</u>, whereas no difference was observed between chicken <u>XX</u> and chicken <u>AA</u> (Figure S20). When extremely weakly expressed genes (FPKM < 0.01) or even unexpressed genes were taken into account, the expression level of human X chromosome was significantly and specifically underestimated, thus resulting in the decreased X:<u>XX</u> ratio. This also aroused another debate over whether all the genes or merely the expressed genes should be investigated [15, 17, 31, 44] [9–11, 14]. In this study, we revealed that the X-linked genes exhibited higher tissue- and developmental stage-specificity and were more in quiescent states (Figure 4), which resulted in more unexpressed X-linked genes in a certain tissue at a certain development stage. We suggested that the expressed genes under a common modest threshold (FPKM > 1) should be considered for a real “fair comparison” when dosage compensation was tested since only expressed genes needed to be compensated and functioned in cells. Meanwhile, one thing that should be mentioned is that abundant non-coding RNAs exist in mammalian tissues, especially in the brain and testis, and exert important regulatory functions essential for tissues despite not encoding proteins [48–50]. Considering the pronounced expression variability and number discrepancy of non-coding RNAs in the different datasets, we only used protein-coding genes to assess the X: AA and X: XX ratio as previous studies did, which makes an easier comparison between our results and past observations. With the improvement of non-coding RNAs annotation in human and other species and the advance in the knowledge of their functions and interactions with protein-coding genes, they could be a complement to testing Ohno’s hypothesis in the future.

Overall, our comprehensive analysis of X:AA and X:<u>XX</u> ratios across tissues, developmental stages, and species support the hypothesis of X chromosome upregulation to compensate for the Y chromosome degeneration.

## Conclusion

In summary, we systematically tested dosage compensation in five mammalian species and confirmed the upregulation of X chromosome at three expression layers across multiple tissues. Combining developmental transcriptome data from organogenesis to adulthood, we observed a dynamic spatial-temporal X:AA ratio but stable dosage compensation (X:XX ratio). Finally, we revealed the difference between X chromosome and autosomes in the tissue-/stage-specificity and underlying epigenetic regulation, which led to the discrepancy of these two expression ratios. Overall, our work supported Ohno’s hypothesis and unified the notion of balanced expression within a genome in most organisms.

## Materials and methods

### Expression analysis

Developmental transcriptome data of human, mouse, and chicken were downloaded from EBI ArrayExpress (www.ebi.ac.uk/arrayexpress) under accession number E-MTAB-6814, E-MTAB-6798, and E-MTAB-6769, respectively. All data generated or analysed during this study are included in Supplemental Table S2. The datasets covered the developmental stages from organogenesis to adulthood across seven major tissues (brain, cerebellum, heart, kidney, liver, ovary, and testis) [20]. We used snakePipes (v1.3.0) [51], a workflow package, for processing high-throughput data, to estimate the gene expression level across tissues in the three species. In RNA-seq analysis, we used SnakePipes to integrate STAR (v2.6.1) and featureCounts (v2.0.0) for mapping reads and for quantifying uniquely mapped reads, respectively. We measured gene expressions with FPKM (Fragments Per Kilobase Million) based on read counts with only protein-coding genes considered. There were 19,694, 21,297 protein-coding genes in human and mouse. Detailed information and resource for public data used in this study were presented in Supplemental Table S2. We directly downloaded the gene expression matrix of rhesus, rat, rabbit, and opossum under accession number E-MTAB-6813, E-MTAB-6811, E-MTAB-6782, and E-MTAB-6833, respectively. To examine the tissue distribution of genes, we assigned a gene to a tissue where this gene showed maximum expression level or protein abundance.

### Tissue specificity and developmental stage specificity

A tau value was used to measure the tissue specificity of genes [52]. For gene A, we defined its mean FPKM value throughout all developmental stages in a certain tissue as gene A’s expression level in this tissue.

The genes with FPKM values of 0 in all tissues were excluded in the analysis. The tau value was calculated in the following formula.

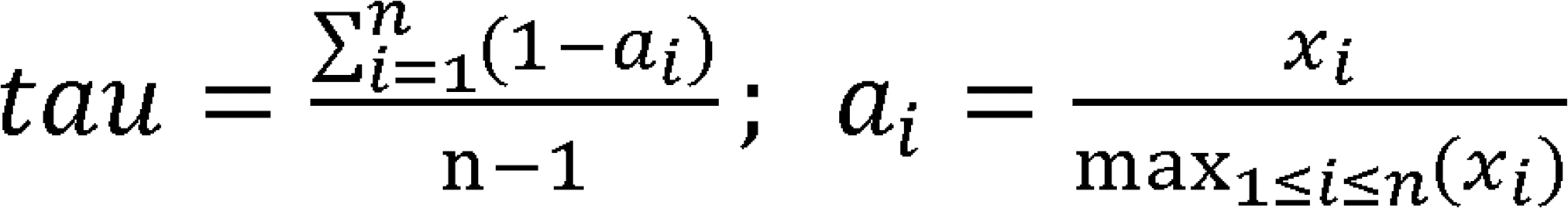

Where the xi means the expression level of gene x in tissue i. The tau value ranged from 0 to 1, and 0 indicated “broadly expressed”, and 1 represented “highly specific”. The same formula was applied to calculate the developmental stage specificity of all genes in each tissue. It should be noted that since the different tissues covered different numbers of sampling stage, the tissue specificity (tau) was not suitable for across- tissue comparison.

### X:AA expression ratio calculation

The ratio of mean expression level of X-linked genes to that of autosomal genes was (X:AA ratio) was calculated, and only genes whose FPKM > 1 were considered. For expression with error, we randomly sampled the genes size-matched with those of X chromosome from autosomes 1000 times, and calculated the expression ratio of X to sampled A. The error bar represented 90% confidence interval.

### Identification of stage-correlated genes

In each tissue, a gene had a series of expression values corresponding to different development stages. We computed the Spearman’s rank correlation coefficient between gene expression values and development stages (from young to old), as previously reported [21]. Stage-positively or negatively-correlated genes meant the genes which showed increased or decreased expressions throughout development stages in a certain tissue, respectively. GO analysis was conducted using clusterProfiler [53].

### Atlas of X:AA ratios in mammals

We combined RNA-seq data of 32 human adult tissues with GTEx data [22]. If there were both RNA-seq data and GTEx data for a certain tissue, the mean value of these two data was defined as X:AA ratio of this tissue. We integrated RNA-seq data of 17 mouse tissues into ENCODE data to calculate the X:AA ratio. We used developmental transcriptome data to estimate the X:AA ratio in rhesus, rat, and rabbit. The visualization was realized using R package gganatogram [54].

### Expression ratio between X and <u>XX</u>

We defined the expression ratio of human X-linked genes to outgroup species (chicken, platypus, and opossum) orthologous genes as X:<u>XX</u> ratio, and that of human autosomal genes to outgroup species orthologous genes as AA:<u>AA</u> ratio. The orthologous gene information was downloaded from ENSEMBL biomart [55]. Since chicken chromosome 1 and 4 are orthologous to human X chromosome [56], 325 human X-linked genes whose orthologous genes were located on chicken chromosome 1 or 4 were investigated when X:<u>XX</u> ratio was calculated. Then, 11,070 human autosomal genes whose orthologous genes were not located on chicken chromosome Z or W were investigated when AA:<u>AA</u> ratio was calculated. Since there were more unexpressed genes on human X chromosome than on human AA, chicken <u>XX</u>, and chicken <u>AA</u> in a given tissue, we analyzed only the genes whose FPKM > 1 for a really fair comparison, and applied a scaling procedure for appropriate across-species comparison. Briefly, we scaled the expression levels to make the median of orthologous gene expression levels equal between species. After normalizing the median of AA:<u>AA</u> ratios into 1, we computed the X:<u>XX</u> ratios. X:<u>XX</u> ratio median of 0.5 indicated the evolutionarily reduced expression of human X-linked gene and no dosage compensation, while X:<u>XX</u> ratio median of 1 indicated evolutionary maintenance of human X-linked gene expression and dosage compensation existence. The opossum chromosome 4, 7, and X were orthologous to human X chromosome [57]. One previous study demonstrated that the double upregulation of opossum X-linked genes in both male and female at single cell level [58], thus the opossum X chromosome was treated as a pair of autosomes in this study, and we repeated the same analysis in opossum as in chicken. A total of 375 and 12,592 gene pairs were used to compute human:opossum X:<u>XX</u> ratio and AA:<u>AA</u> ratio, respectively. A total of 415and 10,740 gene pairs were utilized to calculate human:platypus X:<u>XX</u> ratio and AA:<u>AA</u> ratio, respectively.

### Epigenomic analysis of human genome

Epigenetic data were downloaded from Roadmap Epigenomics Project (http://egg2.wustl.edu/roadmap/; see Supplemental Table S2).In this study, all consolidated epigenomes were included (n□=□127). Detailed information on 15-state chromHMM model and matched colors was presented in the original Roadmap paper (https://egg2.wustl.edu/roadmap/web_portal/chr_state_learning.html#core_15state), while the 15_Quies state was colored with grey (“#e5e5e5”). In 50-state model, the epigenetic states with only hg19-based coordinate were provided, thus we used UCSC liftover to convert it to hg38-based coordinate [59]. We defined upstream 2kb and downstream 1kb regions of transcription start site (TSS) as promoter regions. Bedtools intersect [60] was utilized to obtain the epigenetic states overlapped with promoters. A promoter which had any overlap (>= 1bp) with a state was regarded as annotated by this state.

## CRediT author statement

**Sheng Hu Qian**: Investigation, Data curation, Formal analysis, Validation, Visualization, Writing - original draft, Writing - review & editing. **Yu-Li Xiong**: Investigation, Visualization. **Lu Chen**: Methodology. **Ying-Jie Geng**: Visualization. **Xiao-Man Tang**: Data Curation. **Zhen-Xia Chen**: Conceptualization, Resources, Writing - original draft, Writing - review & editing, Supervision, Funding acquisition.

## Competing interests

The authors declare that they have no competing interests.

## Acknowledgments

We greatly acknowledge Asifa Akhtar and Yidan Sun from Max Planck Institute of Immunobiology and Epigenetics for valuable comments. We also thank all members in our lab for their helpful discussions. Great gratitude goes to Ping Liu from Huazhong Agriculture University for her work at English editing. This work was supported by the National Natural Science Foundation of China [31871305]; Opening Foundation of State Key Laboratory of Freshwater Ecology and Biotechnology, China [2020FB08]; the Fundamental Research Funds for the Central Universities [2662018PY021, 2662019PY003]; and Huazhong Agricultural University Scientific & Technological Self-innovation Foundation [2016RC011]. Funding for open access charge: National Natural Science Foundation of China.

## Supplementary material

**Figure S1 X:AA ratio across human tissues using GTEx RNA-seq data.**

Error bar indicated 90% confidence interval.

**Figure S2 X:AA ratio across mouse tissues using ENCODE RNA-seq data.**

**Figure S3 Cumulative frequencies of genes at translational level.**

**A**) Cumulative frequencies of X-linked genes and autosomal genes in human. A theoretical curve (dashed red line) was generated by doubling X-linked expression. **B**) Expression distributions of X-linked genes and autosomal genes in mouse. **C**) Cumulative frequencies of X-linked genes and autosomal genes in mouse.

**Figure S4 Comparison of protein abundance between X-linked genes and autosomal genes across human tissues**.

**Figure S5 Dynamics of X:AA expression ratio during human development.**

**A**) X:AA ratio in human cerebellum, **B**) heart, **C**) kidney, and **D**) ovary under various thresholds.

**Figure S6 Dynamics of X:AA expression ratio during mouse development.**

**A**) X:AA ratio in mouse brain, **B**) cerebellum, **C**) heart, **D**) kidney, **E**) liver, **F**) testis, and **G**) ovary.

**Figure S7 Expression correlation of one-to-one orthologous genes between human and outgroup species at the transcriptome level.**

**A)** Expression correlation of orthologous genes in brain between human and chicken, B) between human and platypus, **C**) between human and opossum. **D**) Expression correlation of orthologous genes in liver between human and chicken, **E**) between human and platypus, **F**) between human and opossum. **G**) Expression correlation of orthologous genes in testis between human and chicken, **H**) between human and platypus, **I**) between human and opossum.

**Figure S8 Expression correlation of one-to-one orthologous genes between human and outgroup species at the transcriptome level.**

**A)** Expression correlation of orthologous genes in brain between human and chicken, between human and platypus, **C**) between human and opossum. **D**) Expression correlation of orthologous genes in liver between human and chicken, **E**) between human and platypus, **F**) between human and opossum. **G**) Expression correlation of orthologous genes in testis between human and chicken, **H**) between human and platypus, **I**) between human and opossum.

**Figure S9 Comparison of human X with outgroup species XX across testis development.**

**A**) using chicken as outgroup species. **B**) using opossum as outgroup species.

**Figure S10 Expression patern of the X chromosome and autosomes.**

**A)** Distribution of the organ in which genes showed maximum expression at proteome layer. **B**) Tissue specificity, and **C**) stage specificity of genes across all chromosomes.

**Figure S11 Tissue specificity and Stage specificity across chromosomes.**

**A)** Tissue, and **B**) stage specificity in rhesus. **C**) Tissue, and **D**) stage specificity in rat.

**Figure S12 Tissue specificity and Stage specificity across chromosomes.**

**A**) Tissue and **B**) stage specificity in rabbit, **C**) Tissue, and **D**) stage specificity in opossum.

**Figure S13. Epigenetic chromHMM state acorss chromosomes.**

**A**) Number of bases within promoters annotated with each chromHMM state, summed across all 127 epigenomes. **B**) Percentage of state overlapped with promoter of genes on the X and autosomes. **C**) Proportion of state annotated overlapped with promoter of genes on each chromosome annotated with each epigenetic state, summed across all epigenomes

**Figure S14 Percentage of unexpression genes.**

**A**) gene with RPKM < 1 in mouse. **B**) gene with RPKM < 0 in human, **C**) and in mouse.

**Figure S15 Percentage of unexpression genes (FPKM < 1) across human tissues.**

**Figure S16 Percentage of unexpression genes (FPKM < 1) across mouse tissues.**

**Figure S17 Percentage of unexpression genes (FPKM = 0) across human tissues.**

**Figure S18 Percentage of unexpression genes (FPKM = 0) across mouse tissues.**

**Figure S19 Number of X-linked tissue-specific genes in X:AA~0.5 group and X:AA~1 group.**

The Pancreas, saliva secreting gland, liver, and skeletal muscle were divided into X:AA~0.5 group, the rest tissues shown in (Figure 1A) were divided into X:AA~1 group.

**Figure S20 Expression level of F9 during development.**

**A**) In macaque, **B**) in mouse, **C**) in rat, **D**) in rabbit, **E**) in opossum, **F**) in chicken

**Figure S21 Expression level of F9 across 27 human tissues from NCBI.**

**Figure S22 Expression level of SPANXD across 27 human tissues from NCBI.**

**Figure S23 Comparison of unexpressed genes between human and chicken.**

**A**) Percentage of unexpressed genes (FPKM <1) of human X, human AA, chicken XX, and chicken AA. **B**) Percentage of unexpressed genes (FPKM <0) of human X, human AA, chicken XX, and chicken AA. The three lines of significance analysis from top to bottom represent the comparison between human X and human AA, between chicken X and chicken AA, and between human X and chicken X, respectively.

**Table S1 GO analysis of stage positive (rho > 0.8) & negative (rho < −0.8) genes in brain, liver, and testis**

**Table S2 Information and resource of public data used in study**

## Notes

### Competing Interest Statement

The authors have declared no competing interest.

## References

[1] Graves JA. Evolution of vertebrate sex chromosomes and dosage compensation. Nat Rev Genet 2016;17:33–46.

[2] Vicoso B. Molecular and evolutionary dynamics of animal sex-chromosome turnover. Nat Ecol Evol 2019;3:1632–41.

[3] Lenormand T, Fyon F, Sun E, Roze D. Sex Chromosome Degeneration by Regulatory Evolution. Curr Biol 2020;30:3001–6 e5.

[4] Gartler SM. A brief history of dosage compensation. J Genet 2014;93:591–5.

[5] Heard E, Disteche CM. Dosage compensation in mammals: fine-tuning the expression of the X chromosome. Genes Dev 2006;20:1848–67.

[6] Ohno S. Sex chromosomes and sex-linked genes. 1967.

[7] Nguyen DK, Disteche CM. Dosage compensation of the active X chromosome in mammals. Nat Genet 2006;38:47–53.

[8] Lin H, Gupta V, Vermilyea MD, Falciani F, Lee JT, O’Neill LP, et al. Dosage compensation in the mouse balances up-regulation and silencing of X-linked genes. PLoS Biol 2007;5:e326.

[9] Lin H, Halsall JA, Antczak P, O’Neill LP, Falciani F, Turner BM. Relative overexpression of X-linked genes in mouse embryonic stem cells is consistent with Ohno’s hypothesis. Nat Genet 2011;43:1169–70; author reply 71–2.

[10] Deng X, Hiatt JB, Nguyen DK, Ercan S, Sturgill D, Hillier LW, et al. Evidence for compensatory upregulation of expressed X-linked genes in mammals, Caenorhabditis elegans and Drosophila melanogaster. Nat Genet 2011;43:1179–85.

[11] Kharchenko PV, Xi R, Park PJ. Evidence for dosage compensation between the X chromosome and autosomes in mammals. Nat Genet 2011;43:1167–9; author reply 71–2.

[12] Larsson AJM, Coucoravas C, Sandberg R, Reinius B. X-chromosome upregulation is driven by increased burst frequency. Nat Struct Mol Biol 2019;26:963–9.

[13] Fukuda A, Tanino M, Matoba R, Umezawa A, Akutsu H. Imbalance between the expression dosages of X-chromosome and autosomal genes in mammalian oocytes. Sci Rep 2015;5:14101.

[14] Wang ZY, Leushkin E, Liechti A, Ovchinnikova S, Mossinger K, Bruning T, et al. Transcriptome and translatome co-evolution in mammals. Nature 2020.

[15] Chen X, Zhang J. The X to Autosome Expression Ratio in Haploid and Diploid Human Embryonic Stem Cells. Mol Biol Evol 2016;33:3104–7.

[16] Yang JR, Chen X. Dosage sensitivity of X-linked genes in human embryonic single cells. BMC Genomics 2019;20:42.

[17] Lin F, Xing K, Zhang J, He X. Expression reduction in mammalian X chromosome evolution refutes Ohno’s hypothesis of dosage compensation. Proc Natl Acad Sci U S A 2012;109:11752–7.

[18] Chou SJ, Wang C, Sintupisut N, Niou ZX, Lin CH, Li KC, et al. Analysis of spatial-temporal gene expression patterns reveals dynamics and regionalization in developing mouse brain. Sci Rep 2016;6:19274.

[19] Praggastis SA, Thummel CS. Right time, right place: the temporal regulation of developmental gene expression. Genes Dev 2017;31:847–8.

[20] Cardoso-Moreira M, Halbert J, Valloton D, Velten B, Chen C, Shao Y, et al. Gene expression across mammalian organ development. Nature 2019;571:505–9.

[21] Schaum N, Lehallier B, Hahn O, Palovics R, Hosseinzadeh S, Lee SE, et al. Ageing hallmarks exhibit organ-specific temporal signatures. Nature 2020;583:596–602.

[22] Wang D, Eraslan B, Wieland T, Hallstrom B, Hopf T, Zolg DP, et al. A deep proteome and transcriptome abundance atlas of 29 healthy human tissues. Mol Syst Biol 2019;15:e8503.

[23] Consortium GT. The GTEx Consortium atlas of genetic regulatory effects across human tissues. Science 2020;369:1318–30.

[24] Pervouchine DD, Djebali S, Breschi A, Davis CA, Barja PP, Dobin A, et al. Enhanced transcriptome maps from multiple mouse tissues reveal evolutionary constraint in gene expression. Nat Commun 2015;6:5903.

[25] Wilhelm M, Schlegl J, Hahne H, Gholami AM, Lieberenz M, Savitski MM, et al. Mass-spectrometry-based draft of the human proteome. Nature 2014;509:582–7.

[26] Eraslan B, Wang D, Gusic M, Prokisch H, Hallstrom BM, Uhlen M, et al. Quantification and discovery of sequence determinants of protein-per-mRNA amount in 29 human tissues. Mol Syst Biol 2019;15:e8513.

[27] Wang ZY, Leushkin E, Liechti A, Ovchinnikova S, Mossinger K, Bruning T, et al. Transcriptome and translatome co-evolution in mammals. Nature 2020;588:642–7.

[28] Brar GA, Weissman JS. Ribosome profiling reveals the what, when, where and how of protein synthesis. Nat Rev Mol Cell Biol 2015;16:651–64.

[29] Artieri CG, Fraser HB. Evolution at two levels of gene expression in yeast. Genome Res 2014;24:411–21.

[30] Floc’hlay S, Wong E, Zhao B, Viales RR, Thomas-Chollier M, Thieffry D, et al. Cis-acting variation is common across regulatory layers but is often buffered during embryonic development. Genome Res 2020.

[31] Chen J, Wang M, He X, Yang J-R, Chen X. The Evolution of Sex Chromosome Dosage Compensation in Animals. J Genet Genomics 2020:2020.07.04.187476.

[32] Sangrithi MN, Turner JMA. Mammalian X Chromosome Dosage Compensation: Perspectives From the Germ Line. Bioessays 2018;40:e1800024.

[33] Wagner M, Yoshihara M, Douagi I, Damdimopoulos A, Panula S, Petropoulos S, et al. Single-cell analysis of human ovarian cortex identifies distinct cell populations but no oogonial stem cells. Nat Commun 2020;11:1147.

[34] Sarropoulos I, Marin R, Cardoso-Moreira M, Kaessmann H. Developmental dynamics of lncRNAs across mammalian organs and species. Nature 2019;571:510–4.

[35] Emerson JJ, Kaessmann H, Betran E, Long M. Extensive gene traffic on the mammalian X chromosome. Science 2004;303:537–40.

[36] Gurbich TA, Bachtrog D. Gene content evolution on the X chromosome. Curr Opin Genet Dev 2008;18:493–8.

[37] Ellegren H. Sex-chromosome evolution: recent progress and the influence of male and female heterogamety. Nat Rev Genet 2011;12:157–66.

[38] Deng X, Berletch JB, Nguyen DK, Disteche CM. X chromosome regulation: diverse patterns in development, tissues and disease. Nat Rev Genet 2014;15:367–78.

[39] Raznahan A, Disteche CM. X-chromosome regulation and sex differences in brain anatomy. Neurosci Biobehav Rev 2020;120:28–47.

[40] Pehrsson EC, Choudhary MNK, Sundaram V, Wang T. The epigenomic landscape of transposable elements across normal human development and anatomy. Nat Commun 2019;10:5640.

[41] Satterlee JS, Chadwick LH, Tyson FL, McAllister K, Beaver J, Birnbaum L, et al. The NIH Common Fund/Roadmap Epigenomics Program: Successes of a comprehensive consortium. Sci Adv 2019;5:eaaw6507.

[42] Belli M, Shimasaki S. Molecular Aspects and Clinical Relevance of GDF9 and BMP15 in Ovarian Function. Vitam Horm 2018;107:317–48.

[43] Lachmann A, Torre D, Keenan AB, Jagodnik KM, Lee HJ, Wang L, et al. Massive mining of publicly available RNA-seq data from human and mouse. Nat Commun 2018;9:1366.

[44] Xiong Y, Chen X, Chen Z, Wang X, Shi S, Wang X, et al. RNA sequencing shows no dosage compensation of the active X-chromosome. Nat Genet 2010;42:1043–7.

[45] Julien P, Brawand D, Soumillon M, Necsulea A, Liechti A, Schutz F, et al. Mechanisms and evolutionary patterns of mammalian and avian dosage compensation. PLoS Biol 2012;10:e1001328.

[46] Chen X, Zhang J. No X-chromosome dosage compensation in human proteomes. Mol Biol Evol 2015;32:1456–60.

[47] Lahnemann D, Koster J, Szczurek E, McCarthy DJ, Hicks SC, Robinson MD, et al. Eleven grand challenges in single-cell data science. Genome Biol 2020;21:31.

[48] Sun YH, Wang A, Song C, Shankar G, Srivastava RK, Au KF, et al. Single-molecule long-read sequencing reveals a conserved intact long RNA profile in sperm. Nat Commun 2021;12:1361.

[49] Necsulea A, Soumillon M, Warnefors M, Liechti A, Daish T, Zeller U, et al. The evolution of lncRNA repertoires and expression patterns in tetrapods. Nature 2014;505:635–40.

[50] Kleaveland B, Shi CY, Stefano J, Bartel DP. A Network of Noncoding Regulatory RNAs Acts in the Mammalian Brain. Cell 2018;174:350–62 e17.

[51] Bhardwaj V, Heyne S, Sikora K, Rabbani L, Rauer M, Kilpert F, et al. snakePipes: facilitating flexible, scalable and integrative epigenomic analysis. Bioinformatics 2019;35:4757–9.

[52] Yanai I, Benjamin H, Shmoish M, Chalifa-Caspi V, Shklar M, Ophir R, et al. Genome-wide midrange transcription profiles reveal expression level relationships in human tissue specification. Bioinformatics 2005;21:650–9.

[53] Yu G, Wang LG, Han Y, He QY. clusterProfiler: an R package for comparing biological themes among gene clusters. OMICS 2012;16:284–7.

[54] Maag JLV. gganatogram: An R package for modular visualisation of anatograms and tissues based on ggplot2. F1000Res 2018;7:1576.

[55] Howe KL, Contreras-Moreira B, De Silva N, Maslen G, Akanni W, Allen J, et al. Ensembl Genomes 2020-enabling non-vertebrate genomic research. Nucleic Acids Res 2020;48:D689–D95.

[56] International Chicken Genome Sequencing C. Sequence and comparative analysis of the chicken genome provide unique perspectives on vertebrate evolution. Nature 2004;432:695–716.

[57] Mikkelsen TS, Wakefield MJ, Aken B, Amemiya CT, Chang JL, Duke S, et al. Genome of the marsupial Monodelphis domestica reveals innovation in non-coding sequences. Nature 2007;447:167–77.

[58] Mahadevaiah SK, Sangrithi MN, Hirota T, Turner JMA. A single-cell transcriptome atlas of marsupial embryogenesis and X inactivation. Nature 2020;586:612–7.

[59] Lee CM, Barber GP, Casper J, Clawson H, Diekhans M, Gonzalez JN, et al. UCSC Genome Browser enters 20th year. Nucleic Acids Res 2020;48:D756–D61.

[60] Quinlan AR, Hall IM. BEDTools: a flexible suite of utilities for comparing genomic features. Bioinformatics 2010;26:841–2.

